# Pancreatic islet α cells regulate microtubule stability in neighboring β cells to tune insulin secretion and induce functional heterogeneity in individual mouse and human islets

**DOI:** 10.1101/2024.10.21.619544

**Authors:** Kung-Hsien Ho, Syed N. Barmaver, Ruiying Hu, Mahircan Yagan, Hamida K. Ahmed, Irina Kaverina, Guoqiang Gu

## Abstract

We have reported that the microtubule (MT) network in β cells attenuates this function by withdrawing insulin secretory granules (ISGs) away from the plasma membrane. Thus, high glucose-induced MT remodeling is required for robust glucose-stimulated insulin secretion (GSIS). We now show that α-cell secreted hormones, Gcg and/or Glp1, regulate the MT stability in β cells. Activating the receptors of Gcg or Glp1 (GcgR or Glp1R) with chemical agonists induces MT destabilization in β ells in the absence of high glucose. In contrast, inhibiting these receptors with antagonists attenuates high glucose-induced MT destabilization. Supporting the significance of this regulation, the MT networks in β cells of islets with higher α/β cell ratio are less stable than those with lower α/β cell ratio. Within each individual islet, β cells that are located close to α cells show faster MTs remodeling upon glucose stimulation than those away. Consequently, islets with higher α/β cell ratio secrete more insulin in response to high glucose and plasma membrane depolarization, which is recapitulated by direct Gcg stimulation. These combined results reveal a new MT-dependent pathway by which α cells, using Gcg and or Glp1-mediated paracrine signaling, tune β-cell secretion. In addition, the different α-β cell ratios in individual islets lead to their heterogeneous secretory responses, which may be important for handling secretory function needs under different physiological conditions.

**Highlights:**

- Gcg sensitizes glucose-induced MT remodeling in mouse and human β cells
- MT density in single islets anti-correlates with α/β cell ratio
- GSIS levels in single islets positively correlate with α/β cell ratio
- Different α/β cell ratio contributes to heterogeneity of single islet GSIS

## Introduction

Secreting sufficient insulin from islet β cells is a prerequisite for preventing diabetes. Two major factors determine the levels of insulin secretion from any β cell, the level of Ca^2+^ influx that triggers secretion and the numbers of insulin secretory granules (ISGs) that are competent for Ca^2+^-induced secretion (1-4). As the main inducer of insulin secretion, glucose regulates both factors. First, glucose metabolism, after their uptake into β cells, boosts the ATP/ADP ratio, leading to Ca^2+^ influx, triggering secretion (5). Second, intermediate glucose metabolites fuel ISG biosynthesis and movement, increasing the pool of releasable ISGs (6).

Besides glucose, both neuronal and hormonal signals can modulate the levels of Ca^2+^ influx and/or the pool of releasable ISGs. For example, acetyl choline from vagal nerve terminals can potentiate insulin secretion by activating muscarinic acetylcholine receptor 3 that enhances cytoplasmic Ca^2+^ levels (7). In parallel, hormones, including Glp1, Gcg, and Sst from the enteroendocrine system or islets themselves, regulate both Ca^2+^ influx and the pool of releasable ISGs via G-protein-coupled receptors (Gcgr, Glp1r, and Sstr). The classical model is that the binding of these hormones to their receptors either activate (Gcgr or Glp1r, via stimulatory G proteins or Gs) or inhibit (Sstr, via inhibitory G proteins, Gi/o) adenylyl cyclase (8-10). These responses alter the levels of cAMP in β cells, which regulates insulin secretion via impacting Ca^2+^ influx or the Ca^2+^ sensitivity of ISGs (11-13). Intriguingly, we found that cAMP can regulates ISG production via promoting microtubule (MT) nucleation at the Golgi apparatus, underscoring the diverse roles of cAMP in β-cell secretory function (14).

MTs are biopolymers made of α-β tubulin dimers. MT ends have distinct dynamics at their plus- and minus-ends. Their general functions include maintaining cell architecture, regulating organelle function, and aiding cargo transport using motor proteins kinesins and dynein, with the former moving from the minus to plus end and the latter in a reverse direction (15,16). In β cells, most MTs are derived from the Golgi and they form a non-directional meshwork, features that allow them to perform both canonical and non-canonical functions (17). In this regard, high glucose treatment stimulates the nucleation of new MTs from the Golgi in a cAMP-dependent pathway (14), eliciting a canonical function to promote new ISG biosynthesis (18). In contrast, the non-directional nature of the β-cell MT meshwork prevents them from serving as tracks for canonical directional vesicular transport (17,19,20). In fact, our mathematical modeling and experimental data suggest that MTs in β cell periphery tend to orientate in parallel to the plasma membrane, which allows withdrawal of ISGs from underneath the plasma membrane to reduce the pool of releasable ISGs (21,22). Consequently, robust GSIS requires the destabilization of existing MTs and remodeling (17,21). These findings underscore the central regulatory roles of MTs in β-cell secretory function and the importance of understanding the regulation of MT destabilization under glucose stimulation.

We recently reported that microtubule-associated protein tau is required for glucose-induced MT destabilization (21). Under resting glucose, tau has low levels of phosphorylation, allowing it to associate with and stabilize MTs in β cells. High glucose treatment induces tau phosphorylation, which attenuates the affinity of tau to the MTs, leading to its disassociation and subsequent MT destabilization (21). Intriguingly, we found that this above process depends on the activities of several kinases such as Gsk3, PKA, PKC, and CDK5, all of which can be activated by cAMP (23-26). These findings, combined with the roles of Gcg/Glp1 in inducing cAMP production in β cells, led to our hypothesis that islet α cells, by virtue of their secretion of Gcg/Glp1 and induction of cAMP in β cells, regulate the stability of MTs. Here, we test this model in both mouse and human donor islets.

## Research Design and Methods

### Animals

Mouse usage followed protocols approved by the Vanderbilt University IACUC for GG/IK. Euthanasia follows procedures recommended by the AAALAC. Wild type CD-1 (ICR) mice were from Charles River (Wilmington, MA). *Gcg*^*Cre*^ mice was a gift from David Jacobson (Vanderbilt). *R26*^*DTR*^ [57BL/6-Gt(ROSA)26Sortm1(HBEGF)Awi/J] is from Jackson Laboratory. Mouse genotyping follows real-time PCR-based protocol, performed by Transnetyx (Memphis, TN).

### Consideration of sex of mice

Both male and female mice were used at 1:1 ratio whenever possible. Results were combined because islets from male and female mice have yield same results.

### Islet isolation and culture

Islet isolation followed published protocols (17). Briefly, the pancreata of 2 to 5 months old mice were perfused with ∼2 ml of 0.5 mg/mL of collagenase IV (Sigma, St. Louis, MI) dissolved in Hank’s Balanced Salt Solution with Ca^2+^/Mg^2+^. The pancreas was digested at 37°C for ∼10 minutes, washed with RPMI 1640 with 11 mM glucose (Gibco, Dublin, Ireland) and 10% fetal bovine serum (FBS) (i.e., complete RPMI media), followed by hand-picking. Islets were allowed to recover in complete RPMI media for overnight and used for down-stream studies.

### Mouse islet GSIS

Islets were equilibrated in KRB with 2.8 mM glucose (G2.8) for one hour at 37 °C. Around 10 Islets were then transferred into individual wells of 12-well plates to start secretion assays. The conditions used are G2.8, G20, and G20K (20 mM glucose plus 30 mM KCl). For each condition, a 45-minute incubation window is used. Total Ins were assayed after ethanol-HCl extraction. ELISA assays of Ins use kit from Alpco. To induce MT destabilization, 10 µg/ml nocodazole (Noc) was used, added at the step of equilibration, and kept throughout the insulin secretion assays.

### Immunofluorescence (IF) and microscopy

Islets was fixed in [4% PFA + 0.1% saponin (MilliporeSigma)] for overnight at 4 °C or 2-hours at room temperature. They were permeabilized in (50% DMSO + 0.2% Triton X100 + 1XPBS) overnight, and used for routine IF staining. Antibodies used are: mouse anti-E-Cadherin (BD), guinea pig anti-insulin (Dako), mouse anti-Gcg (Dako), Rabbit anti-Gcg (MilliporeSigma), Rabbit anti-Sst (Jackson Immuno Research), goat anti-Sst (Dako), and rabbit anti-detyrosinated tubulin (DeTyr-tubulin) (MilliporeSigma) All secondary antibodies are from Jackson Immuno Research or from Thermo Fisher Scientific. Primary antibodies use 1:100 to 1:2000 dilutions, depending on the estimated number of β cells in each assay. Secondary antibodies use 1:200 to 1:1000 dilutions. Image-capture uses Nikon Eclipse A1R or FV1000 laser scanning confocal microscope with 40X or 100X objectives.

### Human donor islets

Human donor islets were obtained exclusively from IIDP. After receiving, islets were further purified via hand picking, let recover for overnight in CRML-1066 media with 15% FBS, and then used. For secretion assays, islets of similar size were washed and equilibrated with KRB (with G2.8) for 1 hour in 37 °C. Single islets were then transferred into each well of round-bottomed 96-well plates to start the secretion assay. After 45 minutes, islets were transferred to a new well with 50 micro-litter KRB with G11 for 45 minutes, and then 50 micro-litter KRB with G20K. Islets were then lysed for total hormone assays. Insulin/Gcg assays used similar approaches as for mouse samples with Gcg assayed using kits from Mercodia. Islets from 3-4 donors were used.

### MT dynamic assays

For examining MT stability, mouse/human islets were stimulated with G16.5 and fixed at 0, 15, 30, 60, or 120 minutes post stimulation. Insulin, Gcg, and DeTyr-tubulin were stained with whole-mount IF and confocal images captured. The mean fluorescence intensity of DeTyr-tubulin of individual β cells was measured and its relationship with glucose stimulation time and the distance of the β cell to the closest α cell was analyzed.

### Chemical/agonist/antagonist treatment for MT imaging

Islets were pretreated in complete RMPI media containing 2.8 mM glucose, specific chemical, agonists or antagonist for two hours. retreated islets were split into two groups, one stimulated with 20 mM glucose and the other incubated with 2.8 mM glucose. They were then fixed and used for IF staining. For secretion assays, islets were conditioned in KRB with DMSO (controls) or chemicals for one hour before GSIS assays as outlined above. Concentrations used are: Liraglutide: 100 nano-mole (nM); Gcg, 100 nM, Exendin 9, 1 µM, Adomeglivant, 20 µM. For all these chemicals, 2000X stocks were made and kept as frozen stocks, which were then unfrozen and diluted to 1X when need.

### Sample size and statistical analysis

For all mouse assays, at least two batches of studies using different animal or donor islets were conducted, done on different days. Each batch will include 2-4 mice of each sex. Within each batch, at least 2 technical repeats were included, with data points presented separately. For human islets, 3-4 donors were used. Statistical analyses use student t-test, one-way ANOVA, two-way ANOVA, or Pearson’s correlation analysis, depending on the type of study. A p-value ≤ 0.05 was considered significant.

### Resource Availability

Reasonable reagent request will be fulfilled by Guoqiang Gu.

## Results

### GcgR/Glp1R-related signals facilitate glucose-induced MT destabilization in β cells

We tested whether manipulating Glp1R or GcgR activity affects MT stability in β cells. Mouse islets were treated with 2.8 mM glucose (G2.8, basal glucose) or G20 (stimulatory glucose) in the presence of either agonists or antagonists for these two receptors (Fig. 1A). MT stability was assayed via monitoring the levels of DeTyr-tubulin, a well-accepted marker for long-lived microtubule (21,27).

**Fig. 1.**
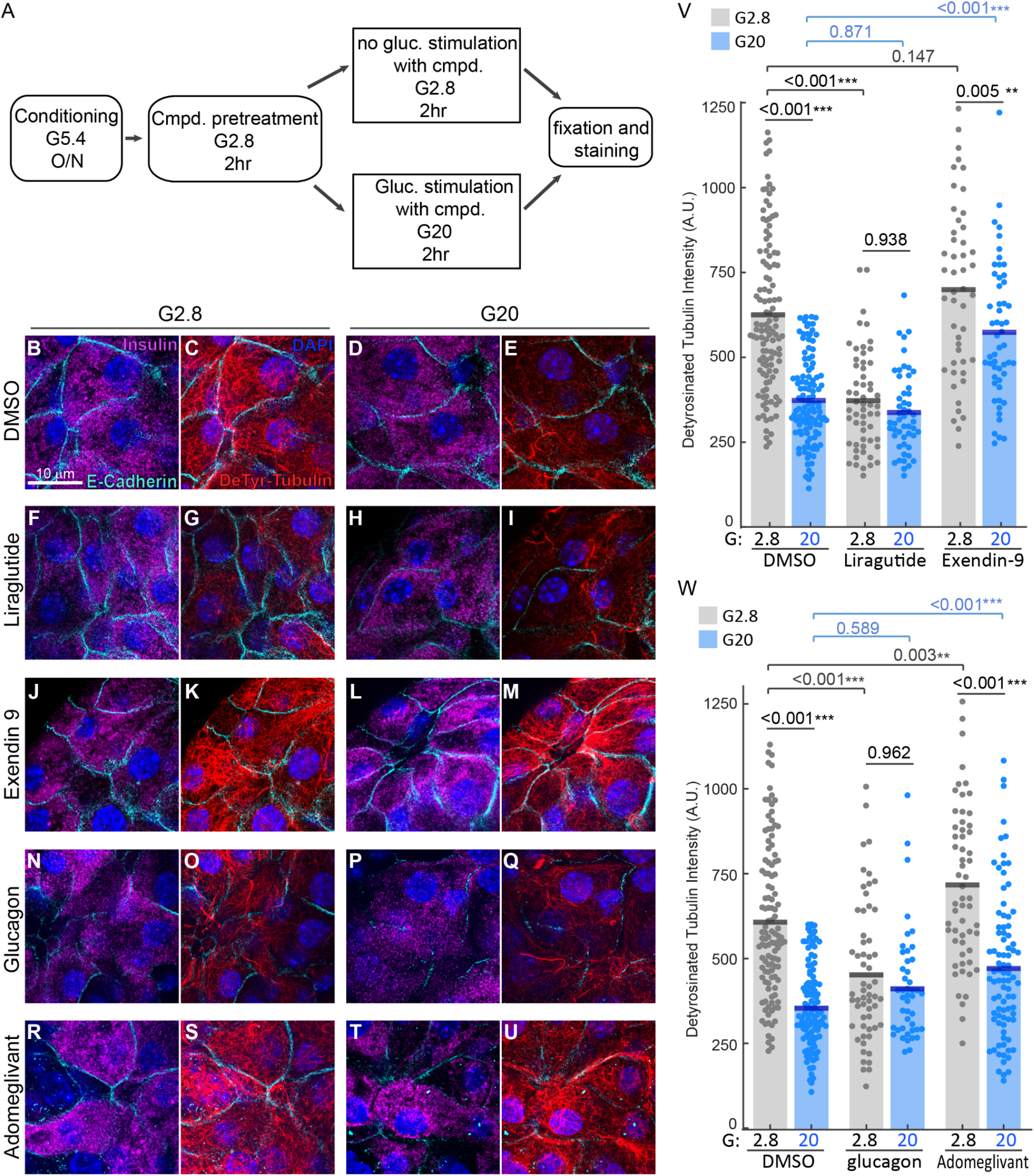
Gcg and/or Glp1 signaling regulates high glucose-induced MT disassembly in β cells. (A) A scheme showing the experimental design. Isolated mouse islets were pre-treated with compounds in media containing 2.8 mM glucose for two hours. Glucose was elevated to 20 mM or stayed in 2.8 mM and incubated for two hours before fixation. (B-U) IF images of detyrosinated (DeTyr) tubulin, E-Cadherin, and insulin in mouse islets cells incubated with 2.8 mM glucose or 20 mM glucose plus 0.05% DMSO (B-E), 100 nM Liraglutide (F-I), 1 *μ*M Exendin-9 (J-M), 100 nM glucagon (N-Q), or 20 *μ*M Adomeglivant (R-U). Blue, DAPI; magenta, insulin; cyan, E-Cadherin; red, DeTyr tubulin. (V-W) Quantification of DeTyr tubulin immunofluorescence intensity in islets β cells using artificial unit (a.u.). Dots represent the mean DeTyr tubulin intensity in the cytoplasm of individual β cells. Bars represent average from three independent repeats (animals). P-values are labeled on the chart (two-way ANOVA with Tukey’s multiple comparison test). ** p<0.01, *** p<0.001.

Mock treated β cells displayed substantially less stable MTs with G20 than those treated with G2.8 (Fig. 1B-E, V). Liraglutide, Glp1R agonist, destabilizes MTs at G2.8 (Fig. 1F-I, V). In contrast, Exendin-9, a Glp1R antagonist, significantly reduces but does not abolish G20-induced MT disassembly (Fig. 1J-M). Similar results were observed when GcgR signaling was activated using Gcg (agonist) or inhibited by Adomeglivant (antagonist) (Fig. 1N-W). Briefly, Gcg significantly destabilizes the MTs in β cells at basal glucose levels. In contrast, Adomeglivant attenuates, but does not eliminate, G20-induced MT destabilization (Fig. 1W). These findings suggest that Gcg and/or Glp1 can directly regulate the MT stability in β cells, leading to different levels of high glucose-induced MT destabilization. We next tested whether α-cell abundance in individual islets impacts the stability of MTs, with the hypothesis that higher levels of GcgR/Glp1R signaling can be induced by higher α/β cell ratio.

### Mouse islets with higher α/β cell ratio have less stable MT in their β cells

Two-month-old *Rosa*^*DTR*^; *Ccg*^*Cre*^ mice were injected with diphtheria toxin (DT). In these mice, the diphtheria toxin (DT) receptor (DTR) is exclusively expressed in Gcg-expressing cells. An injection of DT is expected to ablate a large portion of the α cells (noted as DTRΔ). Three months after DT injection, we detected a reduction of ∼ 70% of α cells in islets with DT injection (assayed by Gcg content in the pancreas. Also compare Fig. 2C and 2G). At the same stage, the MTs are significantly more stable in β cells of DT-injected mice compared with mock-injected mice (Fig. 2A-I). Intriguingly, there is a significant anti-correlation between the abundance of α cells per unit β-cell areas vs MT stability in both the mock- and DT-injected mice (Fig. 2J-K). Note that we did not detect changes of MT stability in the primary cilia of β cells with or without DTRΔ, supporting the specific effects of α cells on cytoplasmic MTs (Fig. 2L).

**Fig. 2.**
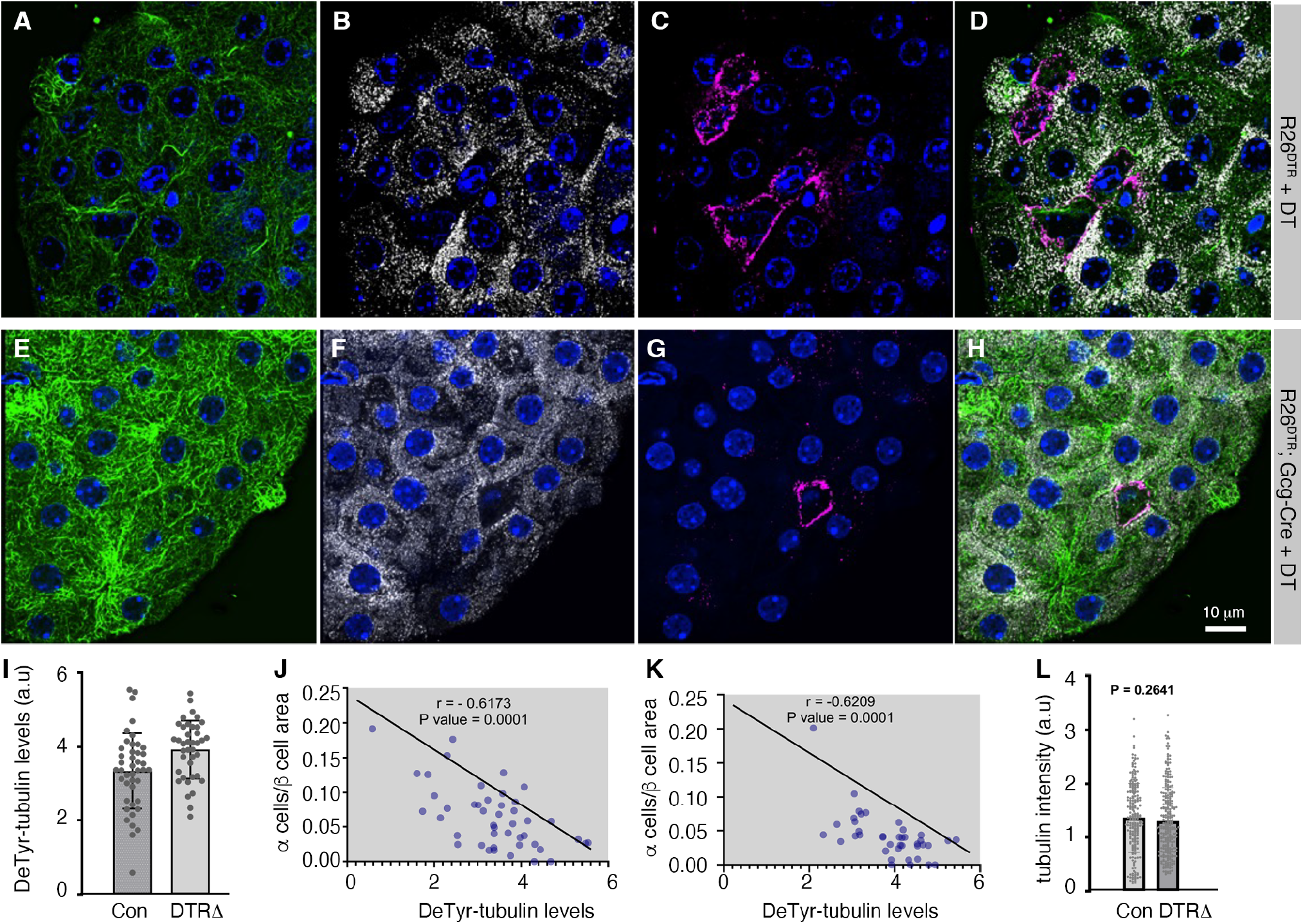
DeTyr-tubulin density anti-correlates with the α/β-cell ratio in mouse islets. (A-H) IF images showing the levels of DeTyr-tubulin (A, E) in β cells without (A-D, control *R26*^*DTR*^ mice) or with DT-induced α cell ablation (E-H, *R26*^*DTR*^; *Gcg-Cre*, noted as DTRΔ). Insulin (B, F) and Gcg (C, G) staining were used to identify the α and β cells, with merged images shown (D, H). Note that all mice received an injection of 20 ng DT each. (I) Quantification of DeTyr-tubulin intensity in islets of control and DTRΔ animals. (J-K) Scatter plots showing a negative correlation between the mean DeTyr-tubulin intensity in β cells and the relative abundance of α cells in control (J) and DTRΔ conditions (K). (L) The DeTyr-tubulin intensity in cilia with or without DT injection. P-values in J and K are from Pearson correlation coefficient test, tailed with CI of 0.95. P-value in L is from t-test, with two-tailed type 2 error.

### MT stability in β cells is influenced by their proximity to α cells

We next examined whether α cells regulate β-cell MT stability using paracrine-like signaling via Gcg/Glp1 diffusion. If so, we expect that the β cells that are closer to α cells will have less stable MTs than those farther away. MT stability in β cells, grouped based on how far they are from any α cell in 3 dimensions, was quantified using confocal image stacks. DeTyr-tubulin content in β cells directly neighboring α cells was compared to β cells separated from α cells by one or two β cell layers. We found that MT stability in β cells of isolated islets significantly anti-correlate with their distance from α cells (Fig. 3), supporting a model that α cells use paracrine-like signaling to destabilize MTs in β cells. These findings, together with the higher GSIS in α-cell enriched islets, led us to directly test whether this α-cell regulated MT destabilization impacts the levels of insulin secretion.

**Fig. 3.**
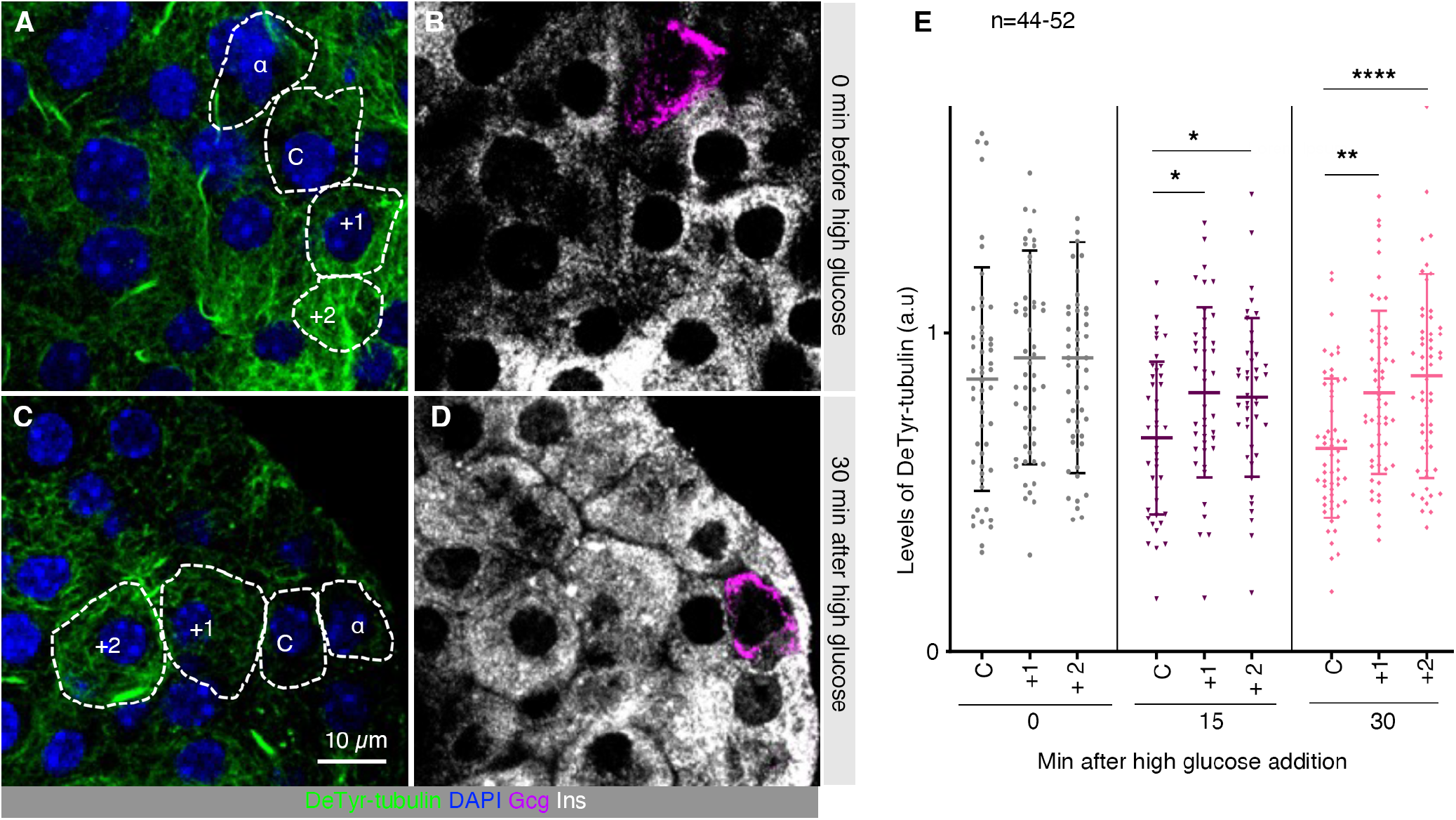
MTs in mouse β cells that are closer to α cells are more sensitive to glucose-induced disassembly. (A, B) Levels of DeTyr-tubulin in mouse β cells before stimulation with high glucose. (C, D) Microtubule stability in β cells with 30 minutes high glucose treatment. Note that a few β cells that directly contact (C), one-cell away (+1), and two-cells away (+2) from an α cell (α) were circled. (E) Quantification of MT stability according to their distance from α cells with 0, 15, or 30 mins treatment by high glucose (G20). Two batches of islets were used, with islets from different mice on different days. *: p<0.05, **: p<0.01, ****: p<0.0001, all calculated with t-test, unpaired type 2 errors.

### Gcg-driven signal(s) and MT destabilization regulate the release of a same pool of ISGs

Insulin secretion was examined with a combination of Gcg stimulation and Noc-induced MT destabilization at basal (G2.8) and stimulatory (G20) glucose conditions (Fig. 4A). We also included a step of stimulation by G20 plus 30 mM KCl (G20K), which will give a maximum stimulation with glucose metabolism and Ca^2+^ influx, revealing the size of releasable ISG pool (Fig. 4A). For this analysis, we utilized islets exclusively from the dorsal half of the mouse pancreas, which contains islets with lower proportions of α cells and having more pronounced response to Gcg-induced insulin secretion (28).

**Fig. 4.**
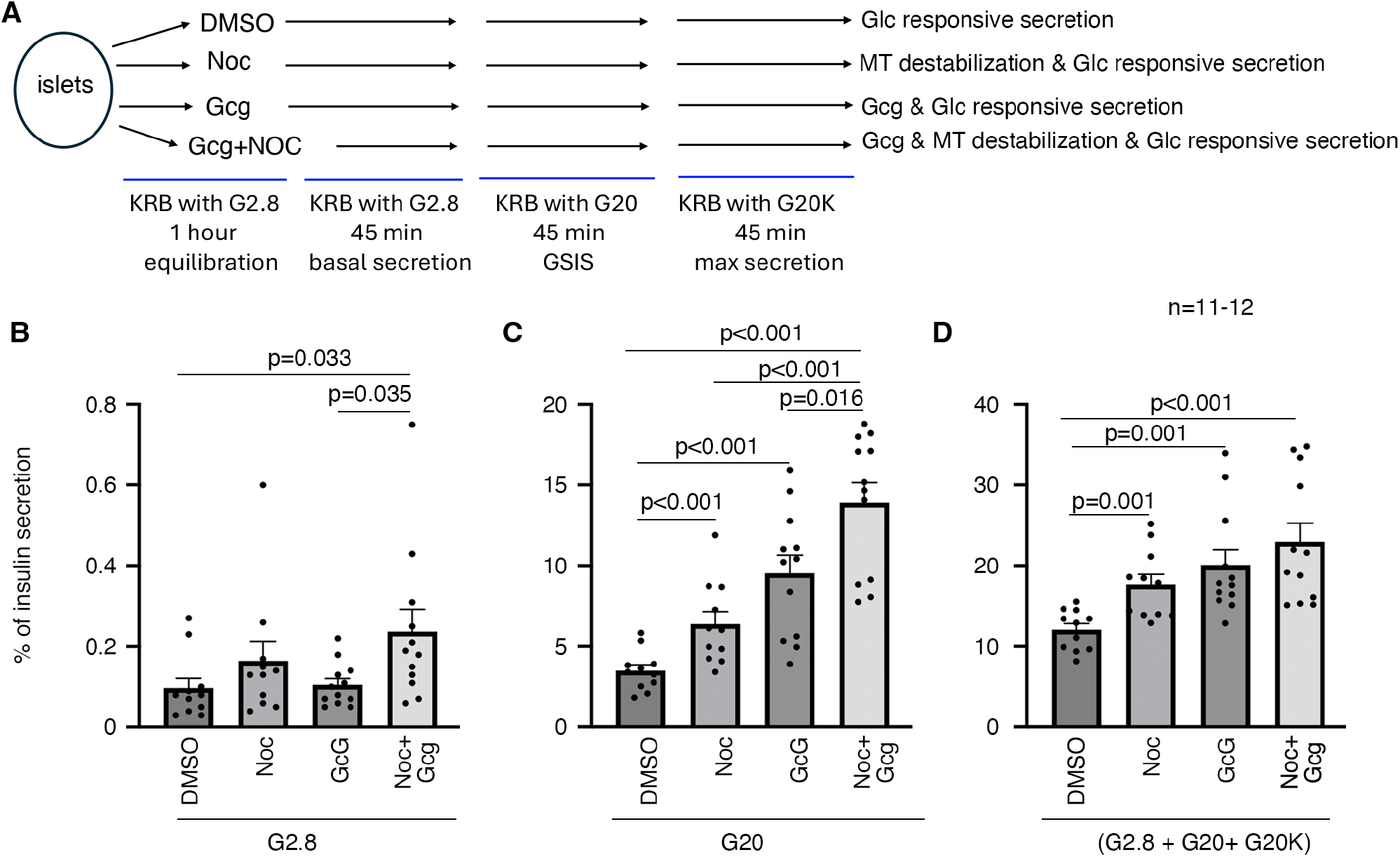
Gcg and MTs regulate an overlapping pool of ISGs for secretion. (A) A scheme showing the experimental design and result interpretation in this study. (B-D) Insulin secretion that are induced by G2.8 alone (B), G20 alone (C), and all conditions (G2.8 + G20 + G20K) (D), with islets pre-treated by different conditions. Presented are (mean + SEM). P-values are calculated with t-test, two-tailed type two error. Only p values that are smaller than 0.050 are presented.

At G2.8, islets treated with Gcg or Noc singly secrete similar levels of insulin as mock-treated islet controls, while islets co-treated with Gcg and nocodazle secrete significantly more (Fig. 4B). At G20, either Gcg or Noc alone significantly boost insulin secretion compared to the control, while co-treatment of both Gcg and nocodazole further enhances secretion (Fig. 4C). When the total releasable insulin was compared (including those induced by G2.8, G20, and G20K), we found that Noc, Gcg, or Noc plus Gcg all induced more insulin secretion than the mock samples (Fig. 4D). However, there is no significant increase in Noc plus Gcg co-treated islets than the Gcg-treated sample (p=0.344). These findings suggest that in the presence of high levels of Ca^2+^ influx, Gcg and MT-destabilization mobilize the same pool of ISGs for secretion. In other words, these data support a model that Gcg induces MT-destabilization in β cells to regulate ISG release. We next examined whether this mechanism is conserved in human islets.

### The α/β cell ratio positively correlates with insulin secretion in human islets

Unlike rodent pancreas where islets in different part of the pancreas have different α/β cell ratio (28), human islets in the head, body, and tail regions of the pancreas have similar proportions of each islet cell type (29). This prevented the pre-separation of islets to different subpopulations based on their α/β cell ratio without terminal antibody staining. We therefore directly assayed insulin secretion in single islets. We focused on islets that are smaller than 300 micrometer in diameter to exclude those with dead cells induced by hypoxia in the core upon receiving.

Proportions of the three major islet cell types, α, β, and δ, were first determined in single islets of four human donors via IF staining assays. Individual human islet showed a large degree of variation, with α/β-cell ratio ranging from 0.00 to 0.84 (Fig. 5A-D). Similarly, the δ to β-cell ratio also showed variation between 0.00 to 0.77 (Fig 5E). Note that there is also a positive correlation between the ratios of α/β and δ/β cells. The functional significance of this finding is not pursued in this study.

**Fig. 5.**
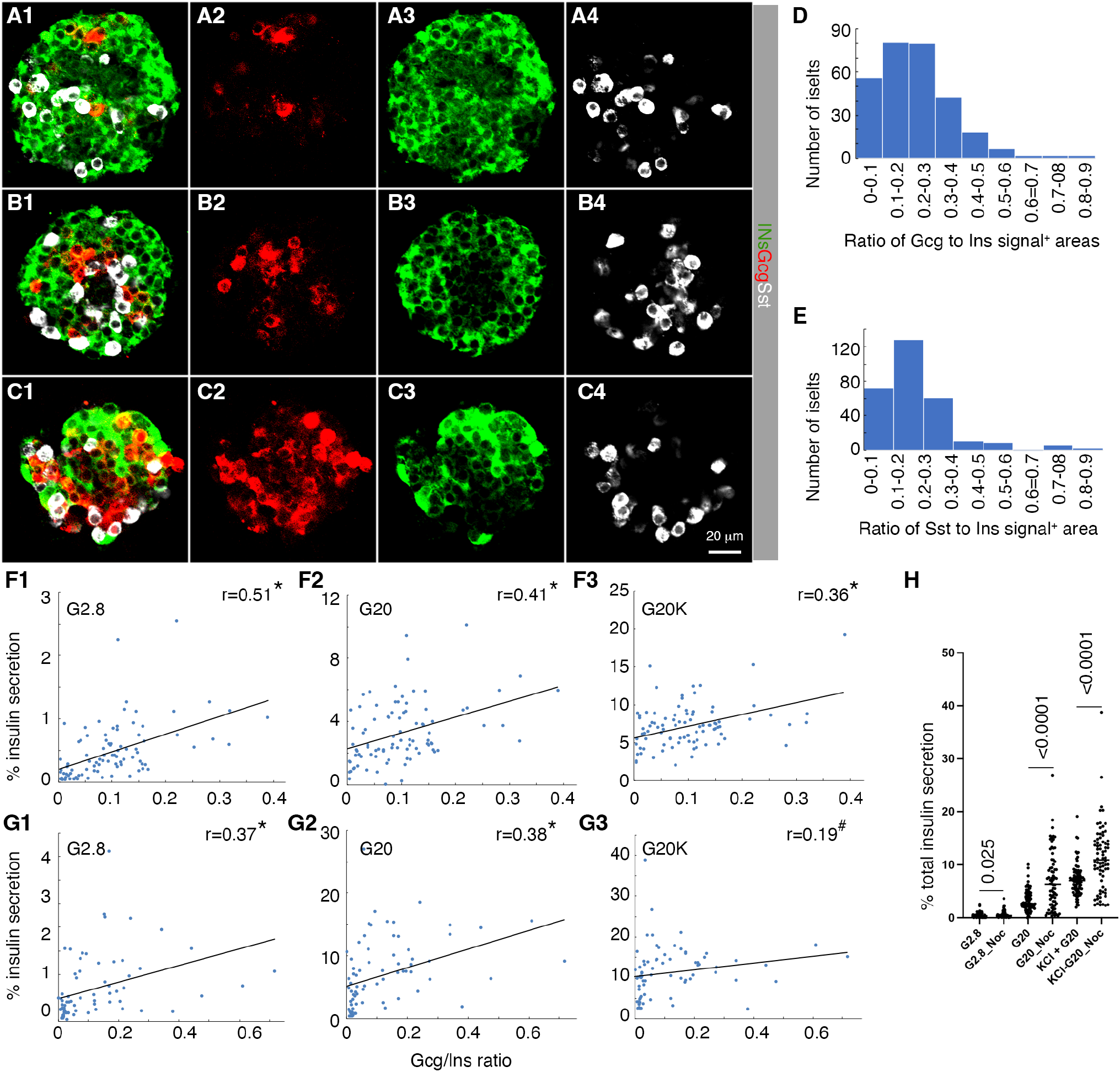
Human islets are heterogeneous in their cell type composition, with α/β cell ratio positively correlates with insulin secretion. (A-C) IF images of three individual human islets with low (A), medium (B), and high (C) α/β cell ratios. Four panels are shown for each islet section, numbered 1 – 4 to note that they are of a same field of view. (D, E) Histograms of islet distribution with different α/β-cell ratios. (F) Correlation of insulin secretion with α-β cell ratio. Sub-panels 1-3 are induced by G2.8, G20, and G20K, respectively. (G) Correlation of insulin secretion in individual human islets after Noc treatment, similar as in F. (G) The Noc-induced insulin secretion in the presence of different stimuli.

Insulin secretion at G2.8, G11, and G20K was assayed in 90 single islets from three donors. A high α/β cell ratio in individual islet is correlated to high insulin secretion at G2.8, G11, and G20K (Fig. 5F). Noc treatment, which destabilizes MTs, does not eliminate the positive correlation between α/β cell ratio and insulin secretion under G2.8 or G11; yet this treatment eliminates this correlation between α/β cell ratio and G20K-induced secretion (Fig. 5H3). These results suggest that similar to mouse islets, α cells in human islets also regulate the same pool of ISGs as those by MTs. Note that this positive correlation does not seem to arise due to the lack of Noc-induced secretion, because Noc significantly enhances the secretion at G2.8, G11, and G20K in these human islets (Fig. 5G).

### Human islet α cells regulates MT stability in neighboring β cells in a paracrine manner

We last tested whether α cells regulate the MT stability in their neighboring β cells in primary human islets. Donor islets were treated with high glucose for different duration of time. MT stability in β cells, grouped based on how far they are from any α cell, were quantified (Fig. 6A-D). We found that glucose stimulation induces MT remodeling in β cells that are in direct contact with α cells earlier than those two cells away (Fig. E). These results suggest that like mouse islets, α cells can regulate MT stability in their neighboring β cells using paracrine signals.

**Fig. 6.**
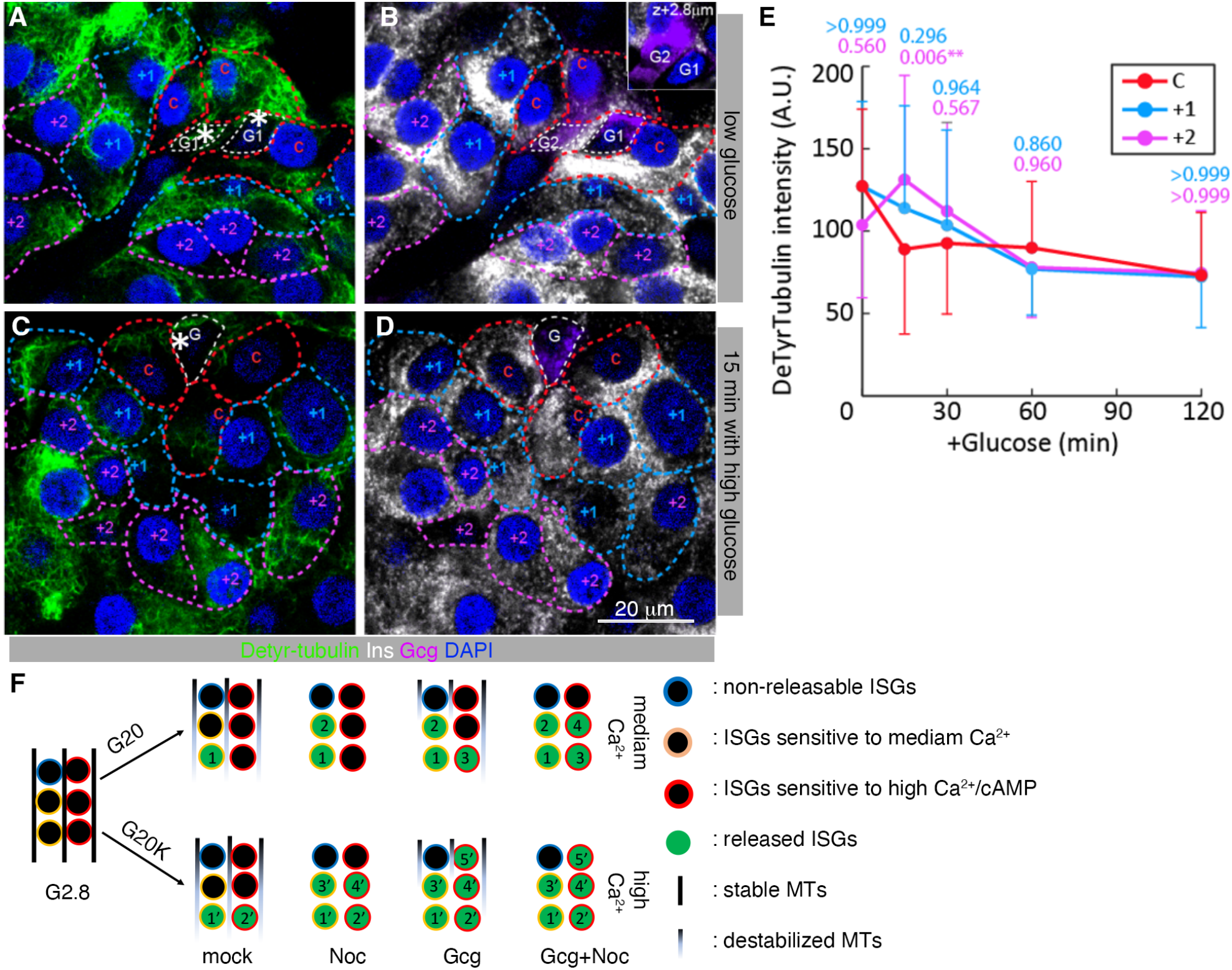
MT stability in human β cells anti-correlates their distance from α cells. (A-D) IF images of DeTyr-tubulin, glucagon, and insulin in human islets cells. Islets were conditioned in media with 5.4 mM glucose overnight (A-B) and stimulated with 16.5 mM glucose for 15 minutes (C-D). Inset in panel B is cropped from the image at the same x-y position but 2.8 micron deeper in the z-axis in the same magnification to show α-cell G2 that is underneath the surrounding β-cells and α-cell G1. Asterisks and G mark the position of α-cells (delineated by white dashed lines). “C” indicates β-cells (delineated by red dashed lines) in direct contact with α-cells. “+ 1” indicates β-cells (delineated by cyan dashed lines) that have one other β-cell between it and the closest α-cell. “+ 2” indicate β-cells (delineated by magenta dashed line) that have two other β-cells between it and the closest α-cell. (E) Quantification of DeTyr-tubulin IF intensity in β-cells stimulated by 16.5 mM glucose for 0, 15, 30, 60, or 120 minutes. Dots represent mean from three replicates (donors). Error bars represent standard deviations. P-values compared “C” and “+1” were labeled in cyan; P-values compared “C” and “+2” were labeled in magenta. ** p<0.01, (two-way ANOVA with Tukey’s multiple comparison test). (F) A proposed model to explain how ISGs in β cells respond to different types of stimulation. Under G20, medium levels of Ca^2+^ influx and partial MT destabilization were induced, allowing the release of a portion of the ISGs with high sensitivity to Ca^2+^ (ISG marked with 1). A complete MT destabilization with Noc allows more secretion (2). Gcg presence not only induces more MT destabilization, but also increased the sensitivity of ISGs to Ca^2+^, boosting the secretion (3, and 4 if Noc is used). Under G20K, high levels of Ca^2+^ influx is induced, allowing the secretion of ISGs with both low and high Ca^2+^ sensitivity (1’ and 2’). Similarly, Noc and Gcg will boost the secretion (those freed by MT destabilization 3’ and 4’ or those by increased Ca^2+^ sensitivity 5’). Yet co-presence of Noc and Gcg cannot further boost secretion because most of the releasable ISGs are released already in the presence of high Gcg alone.

## Discussion

Gcg and insulin work together to maintain blood glucose within a narrow range that is critical for normal physiology (30). Gcg induces gluconeogenesis in the liver to increase blood glucose, while insulin induces glucose uptake and metabolism in peripheral tissues to reduce it. Thus, low glucose induces Gcg secretion from α cells while high glucose induces insulin from β cells (5,31). Further tuning this counter-regulation, high insulin levels repress Gcg secretion, ensuring efficient glucose clearance (32). Counterintuitively, Gcg enhances insulin secretion, which lessens the risk of over-production of glucose and helps to set the resting blood glucose levels (33). The established view is that Gcg activates Gs in β cells, inducing cAMP to enhance Ca^2+^ influx while also increasing the Ca^2+^-sensitivity of ISGs to boost insulin secretion (11-13). By analyzing how Gcg modulates MT dynamics in this study, we suggest that cAMP also regulates both MT polymerization (14) and disassembly (data presented here) to affect the pools of releasable ISGs, which represents another key mechanism by which α cells exert insulinotropic effects.

We recently showed that in islet β cells, MTs have a non-canonical function to withdraw ISGs from the plasma membrane, repressing secretion (17). Upon glucose stimulation, MTs are partially destabilized in a tau-dependent manner, mobilizing ISG for secretion (21). Now we show that α cells can regulate the MT stability in β cells via secreted Gcg and/or Glp1 to impact insulin secretion. These new findings, in combination with those established by others, support the following model: Gcg/Glp1 will activate cAMP production in β cells; cAMP then enhances insulin secretion by: 1) potentiating the Ca^2+^ influx to enhance insulin secretion (11,13); 2) increasing the sensitivity of ISGs to Ca^2+^ (12); and 3) destabilizing MTs to further increase the pools of releasable ISGs (results from this studies). This model can explain our secretion results in the following way (Fig. 6F): G20 induces a medium Ca^2+^ influx while also partially depolymerizing some MTs, allowing the release of a portion of their ISGs having high Ca^2+^ sensitivity. Further MT destabilization by Noc increases secretion by eliminating the MT-mediated ISG withdrawal. Additional Gcg stimulation, via cAMP production, enhances cytoplasmic Ca^2+^ levels, increasing the Ca^2+^ sensitivity of ISGs, while also further destabilizing the MTs. This results in higher levels of secretion. In the co-presence of Noc and Gcg, MTs are completely destroyed, further boost secretion. In the presence of G20K, high Ca^2+^ influx is induced, either MT destabilization (induced by Noc) or ISG modification (by Gcg) is sufficient to induce near complete secretion of the releasable ISG pool.

Our findings have several implications. First, the signals from α cells, mostly relying on Gcg/Glp1, can promote GSIS not only in the short term but also in the long term. To this end, paracrine signaling amongst islet cells has essential roles for balanced hormone secretion and homeostatic function. For example, Ucn3 from β cells can induce the secretion of Sst from δ cells, which in turn represses insulin secretion via Gi-protein coupled receptors in β cells (34). In addition, insulin from β cells can repress Gcg secretion from α cells, help with an efficient clearance of blood glucose, while Gcg enhances GSIS. Our studies suggest that Gcg/Glp1 signaling not only activates GSIS at the short term [likely via modulating Ca^2+^ sensors (e.g., Syt7) via cAMP/PKA (12)], but also modulates β-cell secretion in the long-run via regulating the stability of β-cell MTs.

Second, our findings highlight the functional heterogeneity of endocrine islets. It is now well-established that islet β cells are heterogenous, in their gene expression, Ca^2+^ response, and secretory function (35-39). Such heterogeneity is important for regulated secretion by different types and levels of stimuli (40,41). It also helps to avoid pathological secretion levels, cell dysfunction, and cell death (38,39,42-44). However whether this heterogeneity translates to differences in the secretion of individual islets has not been addressed, although their structural heterogeneity has been documented (45). By correlating the α/β cell ratio in individual islets with their insulin secretion, we establish clear differences in the secretory activities of individual islets. More importantly, we showed that the α/β cell ratio regulates this heterogeneity by modulating the MT network, emphasizing the central regulatory roles of MTs in β-cell insulin secretion.

Third, our findings highlight a new contributor to the disconnection between the physical β-cell mass and the functional β-cell mass. Having a sufficient functional β cell mass is a prerequisite for preventing diabetes (46). Consistent with this model, T2D donors usually have a smaller amount of β cell mass, with a difference of up to 50% (47,48). Yet individual donors, both normal and diabetic patients, can have as high as 6-fold differences in their functional β-cell mass (47). This discrepancy highlights the importance of function in determining β-cell mass, with our studies here showing that α-β cell ratios in islets as a key input for this functional β-cell mass.

There are several interesting issues that remain unanswered. First, we show that Gcg or liraglutides can induce MT destabilization at low glucose. Yet low glucose alone, which induces high levels of Gcg secretion, does not induce observable MT destabilization. It is possible that the α-cell secreted Gcg/Glp1 is not high enough to induce observable destabilization in culture. Alternatively, other unknown signals, induced by high glucose, may be required to induce this stabilization at physiological Gcg/Glp1 levels. Second, we do not know the contribution of δ cells in this MT-regulated insulin secretion process. Sst is known to inhibit GSIS via Gi proteins, lowering the levels of cAMP. Thus, it will be interesting in the future to test if this hormone can affect the glucose-induced MT dynamics in β cells.

## Author contributions

K.H. designed some studies and analyzed hormone expression, MT stability and dynamics in human β cells. S.N.B. did MT stability assays in mouse β cells. R. H. and H.K.A. derived most of the mice needed, helped with islet isolation, and performed immunostainings in some cells. M.Y. did GSIS assays in single human islets and performed whole-mount immune assays. MY helped with IF assays and quantification. IK and GG conceptualized the work and designed most of the studies. All authors helped with writing the manuscript.

## Guarantor Statement

GG is guarantor of the study, with full access to and takes responsibility for the integrity of the data and the accuracy of the data analysis.

## Previous presentation information

The work has not been presented in any places/form.

## References cited

1. West HL, Corbin KL, D’Angelo CV, Donovan LM, Jahan I, Gu G, Nunemaker CS. Postnatal maturation of calcium signaling in islets of Langerhans from neonatal mice. Cell Calcium. 2021;94:102339.

2. Klec C, Ziomek G, Pichler M, Malli R, Graier WF. Calcium Signaling in ss-cell Physiology and Pathology: A Revisit. Int J Mol Sci. 2019;20(24).

3. Blum B, Hrvatin S, Schuetz C, Bonal C, Rezania A, Melton DA. Functional beta-cell maturation is marked by an increased glucose threshold and by expression of urocortin 3. Nat Biotechnol. 2012;30(3):261–264.

4. Huang C, Walker EM, Dadi PK, Hu R, Xu Y, Zhang W, Sanavia T, Mun J, Liu J, Nair GG, Tan HYA, Wang S, Magnuson MA, Stoeckert CJ, Jr., Hebrok M, Gannon M, Han W, Stein R, Jacobson DA, Gu G. Synaptotagmin 4 Regulates Pancreatic beta Cell Maturation by Modulating the Ca(2+) Sensitivity of Insulin Secretion Vesicles. Dev Cell. 2018;45(3):347–361 e345.

5. Deepa Maheshvare M, Raha S, Konig M, Pal D. A pathway model of glucose-stimulated insulin secretion in the pancreatic beta-cell. Front Endocrinol (Lausanne). 2023;14:1185656.

6. Trogden KP, Lee J, Bracey KM, Ho KH, McKinney H, Zhu X, Arpag G, Folland TG, Osipovich AB, Magnuson MA, Zanic M, Gu G, Holmes WR, Kaverina I. Microtubules regulate pancreatic beta-cell heterogeneity via spatiotemporal control of insulin secretion hot spots. Elife. 2021;10.

7. Kong CC, Cheng JD, Wang W. Neurotransmitters regulate beta cells insulin secretion: A neglected factor. World J Clin Cases. 2023;11(28):6670–6679.

8. Dickerson MT, Dadi PK, Zaborska KE, Nakhe AY, Schaub CM, Dobson JR, Wright NM, Lynch JC, Scott CF, Robinson LD, Jacobson DA. G(i/o) protein-coupled receptor inhibition of beta-cell electrical excitability and insulin secretion depends on Na(+)/K(+) ATPase activation. Nat Commun. 2022;13(1):6461.

9. Oliveira de Souza C, Sun X, Oh D. Metabolic Functions of G Protein-Coupled Receptors and beta-Arrestin-Mediated Signaling Pathways in the Pathophysiology of Type 2 Diabetes and Obesity. Front Endocrinol (Lausanne). 2021;12:715877.

10. Mayendraraj A, Rosenkilde MM, Gasbjerg LS. GLP-1 and GIP receptor signaling in beta cells - A review of receptor interactions and co-stimulation. Peptides. 2022;151:170749.

11. Rajan AS, Hill RS, Boyd AE, 3rd. Effect of rise in cAMP levels on Ca2+ influx through voltage-dependent Ca2+ channels in HIT cells. Second-messenger synarchy in beta-cells. Diabetes. 1989;38(7):874–880.

12. Wu B, Wei S, Petersen N, Ali Y, Wang X, Bacaj T, Rorsman P, Hong W, Sudhof TC, Han W. Synaptotagmin-7 phosphorylation mediates GLP-1-dependent potentiation of insulin secretion from beta-cells. Proc Natl Acad Sci U S A. 2015;112(32):9996–10001.

13. Braun M. The somatostatin receptor in human pancreatic beta-cells. Vitam Horm. 2014;95:165–193.

14. Trogden KP, Zhu X, Lee JS, Wright CVE, Gu G, Kaverina I. Regulation of Glucose-Dependent Golgi-Derived Microtubules by cAMP/EPAC2 Promotes Secretory Vesicle Biogenesis in Pancreatic beta Cells. Curr Biol. 2019;29(14):2339–2350 e2335.

15. Barlan K, Gelfand VI. Microtubule-Based Transport and the Distribution, Tethering, and Organization of Organelles. Cold Spring Harb Perspect Biol. 2017;9(5).

16. Logan CM, Menko AS. Microtubules: Evolving roles and critical cellular interactions. Exp Biol Med (Maywood). 2019;244(15):1240–1254.

17. Zhu X, Hu R, Brissova M, Stein RW, Powers AC, Gu G, Kaverina I. Microtubules Negatively Regulate Insulin Secretion in Pancreatic beta Cells. Dev Cell. 2015;34(6):656–668.

18. Schmoranzer J, Simon SM. Role of microtubules in fusion of post-Golgi vesicles to the plasma membrane. Mol Biol Cell. 2003;14(4):1558–1569.

19. Varadi A, Ainscow EK, Allan VJ, Rutter GA. Involvement of conventional kinesin in glucose-stimulated secretory granule movements and exocytosis in clonal pancreatic beta-cells. J Cell Sci. 2002;115(Pt 21):4177–4189.

20. Varadi A, Tsuboi T, Johnson-Cadwell LI, Allan VJ, Rutter GA. Kinesin I and cytoplasmic dynein orchestrate glucose-stimulated insulin-containing vesicle movements in clonal MIN6 beta-cells. Biochem Biophys Res Commun. 2003;311(2):272–282.

21. Ho KH, Yang X, Osipovich AB, Cabrera O, Hayashi ML, Magnuson MA, Gu G, Kaverina I. Glucose Regulates Microtubule Disassembly and the Dose of Insulin Secretion via Tau Phosphorylation. Diabetes. 2020;69(9):1936–1947.

22. Bracey KM, Ho KH, Yampolsky D, Gu G, Kaverina I, Holmes WR. Microtubules Regulate Localization and Availability of Insulin Granules in Pancreatic Beta Cells. Biophys J. 2020;118(1):193–206.

23. Khaled M, Larribere L, Bille K, Aberdam E, Ortonne JP, Ballotti R, Bertolotto C. Glycogen synthase kinase 3beta is activated by cAMP and plays an active role in the regulation of melanogenesis. J Biol Chem. 2002;277(37):33690–33697.

24. Sassone-Corsi P. The cyclic AMP pathway. Cold Spring Harb Perspect Biol. 2012;4(12).

25. Han P, Sonati P, Rubin C, Michaeli T. PDE7A1, a cAMP-specific phosphodiesterase, inhibits cAMP-dependent protein kinase by a direct interaction with C. J Biol Chem. 2006;281(22):15050–15057.

26. Chen MC, Lin H, Hsu FN, Huang PH, Lee GS, Wang PS. Involvement of cAMP in nerve growth factor-triggered p35/Cdk5 activation and differentiation in PC12 cells. Am J Physiol Cell Physiol. 2010;299(2):C516–527.

27. Hu R, Zhu X, Yuan M, Ho KH, Kaverina I, Gu G. Microtubules and Galphao-signaling modulate the preferential secretion of young insulin secretory granules in islet beta cells via independent pathways. PLoS One. 2021;16(7):e0241939.

28. Trimble ER, Halban PA, Wollheim CB, Renold AE. Functional differences between rat islets of ventral and dorsal pancreatic origin. J Clin Invest. 1982;69(2):405–413.

29. Walker JT, Saunders DC, Rai V, Chen HH, Orchard P, Dai C, Pettway YD, Hopkirk AL, Reihsmann CV, Tao Y, Fan S, Shrestha S, Varshney A, Petty LE, Wright JJ, Ventresca C, Agarwala S, Aramandla R, Poffenberger G, Jenkins R, Mei S, Hart NJ, Phillips S, Kang H, Greiner DL, Shultz LD, Bottino R, Liu J, Below JE, Consortium H, Parker SCJ, Powers AC, Brissova M. Genetic risk converges on regulatory networks mediating early type 2 diabetes. Nature. 2023;624(7992):621–629.

30. Roder PV, Wu B, Liu Y, Han W. Pancreatic regulation of glucose homeostasis. Exp Mol Med. 2016;48(3):e219.

31. Zhang J, Zheng Y, Martens L, Pfeiffer AFH. The Regulation and Secretion of Glucagon in Response to Nutrient Composition: Unraveling Their Intricate Mechanisms. Nutrients. 2023;15(18).

32. Cooperberg BA, Cryer PE. Insulin reciprocally regulates glucagon secretion in humans. Diabetes. 2010;59(11):2936–2940.

33. Rodriguez-Diaz R, Molano RD, Weitz JR, Abdulreda MH, Berman DM, Leibiger B, Leibiger IB, Kenyon NS, Ricordi C, Pileggi A, Caicedo A, Berggren PO. Paracrine Interactions within the Pancreatic Islet Determine the Glycemic Set Point. Cell Metab. 2018;27(3):549–558 e544.

34. van der Meulen T, Donaldson CJ, Caceres E, Hunter AE, Cowing-Zitron C, Pound LD, Adams MW, Zembrzycki A, Grove KL, Huising MO. Urocortin3 mediates somatostatin-dependent negative feedback control of insulin secretion. Nat Med. 2015;21(7):769–776.

35. Johnston NR, Mitchell RK, Haythorne E, Pessoa MP, Semplici F, Ferrer J, Piemonti L, Marchetti P, Bugliani M, Bosco D, Berishvili E, Duncanson P, Watkinson M, Broichhagen J, Trauner D, Rutter GA, Hodson DJ. Beta Cell Hubs Dictate Pancreatic Islet Responses to Glucose. Cell Metab. 2016;24(3):389–401.

36. Zhang M, Goforth P, Bertram R, Sherman A, Satin L. The Ca2+ Dynamics of Isolated Mouse β-Cells and Islets: Implications for Mathematical Models. Biophysical Journal. 2003/05/01;84(5).

37. Giordano E, Bosco D, Cirulli V, Meda P. Repeated glucose stimulation reveals distinct and lasting secretion patterns of individual rat pancreatic B cells. The Journal of clinical investigation. 1991 Jun;87(6).

38. Schravendijk CFV, Kiekens R, Pipeleers DG. Pancreatic beta cell heterogeneity in glucose-induced insulin secretion. Journal of Biological Chemistry. 1992/10/25;267(30).

39. Wojtusciszyn A, Armanet M, Morel P, Berney T, Bosco D. Insulin secretion from human beta cells is heterogeneous and dependent on cell-to-cell contacts - PubMed. Diabetologia. 2008 Oct;51(10).

40. Benninger RKP, Dorrell C, Hodson DJ, Rutter GA. The Impact of Pancreatic Beta Cell Heterogeneity on Type 1 Diabetes Pathogenesis. Curr Diab Rep. 2018;18(11):112.

41. Benninger RKP, Kravets V. The physiological role of beta-cell heterogeneity in pancreatic islet function. Nat Rev Endocrinol. 2022;18(1):9–22.

42. Dominguez-Gutierrez G, Xin Y, Gromada J. Heterogeneity of human pancreatic beta-cells. Mol Metab. 2019;27S(Suppl):S7–S14.

43. Wojtusciszyn A, Armanet M, Morel P, Berney T, Bosco D, Wojtusciszyn A, Armanet M, Morel P, Berney T, Bosco D. Insulin secretion from human beta cells is heterogeneous and dependent on cell-to-cell contacts. Diabetologia 2008 51:10. 2008-07-30;51(10).

44. Dominguez-Gutierrez G, Xin Y, Gromada J. Heterogeneity of the Pancreatic Beta Cell. Frontiers in genetics. 03/06/2017;8.

45. Dybala MP, Hara M. Heterogeneity of the Human Pancreatic Islet. Diabetes. 2019;68(6):1230–1239.

46. Weir GC, Gaglia J, Bonner-Weir S. Inadequate beta-cell mass is essential for the pathogenesis of type 2 diabetes. Lancet Diabetes Endocrinol. 2020;8(3):249–256.

47. Rahier J, Guiot Y, Goebbels RM, Sempoux C, Henquin JC. Pancreatic beta-cell mass in European subjects with type 2 diabetes. Diabetes Obes Metab. 2008;10 Suppl 4:32–42.

48. Ritzel RA, Butler AE, Rizza RA, Veldhuis JD, Butler PC. Relationship between beta-cell mass and fasting blood glucose concentration in humans. Diabetes care. 2006;29(3):717–718.

